# Use of cellular FAD autofluorescence as a label-free cellular attribute for the production of chimeric antigen receptor-T cells

**DOI:** 10.1101/2025.01.14.633077

**Authors:** Ka-Wai Cheung, Faris Kairi, Denise Bei Lin Teo, Wei-Xiang Sin, Yie Hou Lee, Michael E. Birnbaum

## Abstract

Chimeric antigen receptor T (CAR-T) cell therapy has become an attractive approach for treating hematological malignancies. However, the accessibility of this therapy is limited by factors such as complex manufacturing process, limited capacity of manufacturing facilities and the requirement of highly skilled workforce for the manual steps of CAR-T cell production. To minimize the manual processes, CAR-T cell manufacturing field is shifting towards closed and automated systems, including analytical tools that offer intermittent monitoring of cells in production. Therefore, label-free technologies for closely monitoring CAR-T cells in closed systems are needed. Here, we evaluate the use of a flow cytometer equipped with a 405nm violet laser for investigating the NADH and FAD autofluorescence in T cells. Our results revealed the increase of NADH and FAD autofluorescence were significantly correlated with the upregulation of T cell activation marker, CD25 and the increase of extracellular lactate in spent media in the first three days after T cell activation. We demonstrate the potential use of FAD for determining the endpoint of CAR-T cell manufacture by establishing a relationship between the rate of change in the mean fluorescence intensity (MFI) of FAD in CAR-T cells and the rate of change in T cell proliferation using a G-Rex bioreactor. Collectively, these findings suggest that autofluorescence, particularly FAD autofluorescence, can serve as a label-free biomarker (cellular attribute) for monitoring T cell activation and expansion during CAR-T cell production. The use of 405nm visible light to substitute the genotoxic UV wavelengths for assessing the NADH and FAD autofluorescence, paves the way to incorporate autofluorescence measurements into closed and automated systems for in-process monitoring of CAR-T cell manufacturing.

## Introduction

Chimeric antigen receptor T (CAR-T) cells are genetically modified T cells which express synthetic receptors on the cell surface for identifying specific tumor antigens and subsequently, eradicate the tumor cells. Since the launch of CAR-T cell products in 2017, CAR-T cell therapy has become an attractive approach for cancer treatment^1,2^. To date, seven US FDA approved CAR-T cell therapies against B cell malignancies including leukemia, lymphoma and multiple myeloma are commercially available on the market^3–5^, with a large number of candidates, targeting both hematological malignancies and solid tumors undergoing different stages of clinical trials^6,7^.

CAR-T cell production starts with the collection of peripheral blood mononuclear cells (PBMCs) through leukapheresis, followed by T cell enrichment. The T cells are then activated, genetically modified via viral or non-viral gene delivery and expanded, typically for 9 to 14 days^3,8–11^. While the general workflow is largely consistent across products and laboratories, the complex manufacturing process, limited capacity of manufacturing facilities and the requirement of highly skilled workforce for the manual steps of CAR-T cell production have limited the accessibility of these therapies for patients^11,12^. Hence, future developments in CAR-T cell manufacturing are focusing on shifting from manual processes to closed and automated systems.

Current closed and automated systems, such as CliniMACS Prodigy and Lonza Cocoon enable key CAR-T cell manufacturing processes (T cell isolation, activation, transduction and expansion) to be carried out on a platform that remains sealed from the environment. However, sample removal for cell analysis during the manufacturing process requires manual handling, even though the procedures utilize sterile tubing welders or aseptic access ports to maintain sterility^12^. To reduce the manual processing and manufacturing interventions, label-free in-line or on-line technologies for closely monitoring the manufacturing processes such as T cell activation and expansion in closed systems are needed.

Nicotinamide adenine dinucleotide reduced (NADH) and flavin adenine dinucleotide (FAD) are endogenous molecules involved in cellular metabolism including glycolysis and oxidative phosphorylation (OXPHOS)^13,14^ and can be excited by wavelength 330-360nm and 360-465nm to emit cellular autofluorescence, respectively^14^. Previous studies have shown that NADH and FAD autofluorescence can serve as biomarkers for T cell activation^15,16^, indicating that these molecules represent potential targets for the development of label-free technologies to monitor CAR-T cell manufacturing processes in closed systems. Lemire, et. al. has revealed the ultraviolet (UV) laser source equipped flow cytometer can be used to measure the NADH and FAD autofluorescence in immune cells^17^. Unfortunately, since UV wavelengths induce genotoxicity in cells, a UV laser source is not well suited for CAR-T cell manufacture. The use of two-photon microscopy enables the NADH and FAD autofluorescence to be induced by less harmful infrared wavelengths^15,16,18^. Although two-photon microscopy can provide information on the T cell metabolism through fluorescence lifetime imaging, limitations of two-photon microscopy, such as imaging depth^19^, the number of cells analyzed per image, and the requirement for cells to settle down to the bottom before image capture^17,18^ render this method unsuitable for the suspension CAR-T cells cultured in a dynamic environment.

In contrast to two-photon microscopy, flow cytometry is commonly used in immune cell profiling, and can assess key attributes of CAR-T therapies (including CD4/8 ratio, CAR expression, memory phenotypes, and state of cell exhaustion)^20,21^. Therefore, flow cytometry is an attractive means of assessing CAR-T cells given its robustness, speed of analysis, and familiarity to cell therapy manufacturers. However, these measurements typically require staining the sample, which is not compatible with real-time monitoring. It has previously been demonstrated that changes in NADH and FAD autofluorescence can be quantified using flow cytometry^17^. In this study, we demonstrate that changes in NADH and FAD autofluorescence can be assessed with a flow cytometer equipped with a 405nm laser to monitor T cell activation. In addition, we demonstrate the potential use of FAD as a cellular attribute for determining the endpoint of CAR-T cell manufacture in a label free manner that could be integrated as an in-line process measurement that does not require genotoxic UV light.

## Materials and Methods

### Production of lentivirus for generating CD19CAR-T cells (CD19TDC)

pMDLg/pRRE, pRSV-Rev and pMD2.G packaging plasmids and anti-CD19 CAR-IRES-EGFP transfer plasmid were used for producing lentivirus for generating CD19TDC. The packaging plasmids were gifts from D. Trono (Addgene plasmid numbers 12251, 12253 and 12259). The details of the transfer plasmids and the methods for generating the transfer plasmid are as described previously^22^. The detailed method for the lentivirus production and titration are as described in a previous publication^21^.

### T cell isolation

Frozen human peripheral blood mononuclear cells (PBMCs) from healthy donors were purchased from STEMCELL Technologies. CD3+ T cells (T cells) were purified from the thawed PBMCs (Purity: >98%) using EasySep Human T Cell Isolation Kit (STEMCELL Technologies).

### Cell preparation for measuring the NADH and FAD autofluorescence after T cell activation

1×10^6^/ml T cells were stimulated with 25ul/ml Immunocult Human CD3/CD28 T cell activator (Immunocult or IC) (STEMCELL Technologies) in a 24-well plate (Thermo Fisher Scientific) in full medium ((AIM-V Medium, Thermo Fisher Scientific) supplemented with 2% human male AB serum (Merck)) in the presence of 100 IU/ml recombinant human IL-2 (Miltenyi Biotec) for 3 days. Unstimulated cells were cultured without Immunocult and IL-2.

### Cell preparation for measuring the longitudinal changes of NADH and FAD autofluorescence in T cells

2×10^6^ T cells were stimulated by incubating with Dynabeads Human T-Expander CD3/CD28 (Dynabeads) (Thermo Fisher Scientific) at a 1:1 cell:bead ratio in 8ml full medium with 100 IU/ml IL-2. To study the effect of cytokines on autofluorescence, Dynabeads stimulated T cells were cultured under the same condition except replacing the IL-2 with a combination of 12.5ng/ml IL-7 and 12.5ng/ml IL-15 (Miltenyi Biotec). To study the effect of activation strength on autofluorescence, 2×10^6^ T cells were stimulated with 25ul/ml or 2.5ul/ml Immunocult in 8ml full medium with IL-2. T cells were expanded in a G-Rex 24-well plate (Wilson Wolf) until day 14 at a constant volume of 8ml per well, with 5ml medium replenished with fresh full medium with IL-2 or a combination of IL-7 and IL-15 daily over the course of 14 days.

### Cell preparation for studying the longitudinal changes of NADH and FAD autofluorescence in non-transduced T cells (NTC) and CD19TDC

2×10^6^ T cells were stimulated by incubating with Dynabeads Human T-Expander CD3/CD28 (Thermo Fisher Scientific) at a 1:1 cell:bead ratio in 2ml full medium with 100 IU/ml IL-2 in a G-Rex 24-well plate. One day after activation, 1ml of cell culture supernatant was removed carefully without disturbing the cells settled in the bottom of the well. For CD19TDC, 1ml of full medium with IL-2 containing diluted lentivirus was subsequently added to the well at a MOI of 5. 1ml of full medium with IL-2 alone was added to NTC. One day after transduction, 6ml of full medium with IL-2 was added to achieve the final culture volume of 8ml per well for both NTC and CD19TDC. The cells were then expanded until day 14 with 5ml medium replenished with fresh full medium with IL-2 daily over the course of 14 days.

### Antibodies and reagents for flow cytometry related experiments

The following fluorochrome-conjugated anti-human antibodies and their relevant isotype controls were used for flow cytometry and were purchased from BioLegend: FITC anti-CD3 (UCHT1), APC anti-CD19 (HIB19), BV510 anti-CD4 (clone OKT4), FITC anti-CD8 (clone HIT8a), PECy7 anti-CD25 (clone BC96), APC anti-CD69 (clone FN50), BV605 anti-CD45RA (clone HI100), PE anti-CCR7 (clone G043H7), APC mouse IgG1,κ isotype (MOPC-21), PECy7 mouse IgG1, κ isotype (MOPC-21), PE Mouse IgG2a,κ isotype (MOPC-173) and BV605 Mouse IgG2b,κ isotype (MPC-11). LIVE/DEAD fixable violet dead cell stain kit (LiveDead-violet), SYTOX-AADvanced Dead cell stain Kit (sytox-AAD) and CountBright^TM^ absolute counting beads were purchased from Thermo Fisher Scientific.

### FACS analysis for measuring the NADH and FAD autofluorescence after T cell activation

Unstimulated and stimulated T cells were retrieved from the 24 well plates followed by washing twice with FACS buffer (PBS supplemented with 2% Fetal Bovine Serum (FBS) (Thermo Fisher Scientific) and 0.1% sodium azide (Merck)). Autofluorescence data were acquired by a CytoFLEX S flow cytometer equipped with a 405nm violet laser, without an UV laser (Beckman Coulter). NADH and FAD autofluorescence were excited by 405nm violet laser and the signals were detected with bandpass filters BP 450/45 and BP 525/40, respectively. Data were analyzed using CytExpert software (Beckman Coulter). An example of the gating strategy for percentage and MFI of FAD is illustrated in Suppl. Figure 1.

### FACS analysis for measuring the longitudinal changes of NADH and FAD autofluorescence and cell count

Two aliquots of 20µl of T cells were retrieved from a well of a G-Rex plate for analysis during medium replenishment daily until day 14. For NTC and CD19TDC, the two aliquots were retrieved from the 2ml culture in a G-Rex plate for the first two days and same procedure was performed as above for T cells since day 3. The first aliquot was used for analyzing the autofluorescence in unstained cells. The cells in the second aliquot were stained with sytox-AAD for dead cell discrimination and used for analyzing the autofluorescence in living cells. For FACS analysis, 330ul FACS buffer were added to each of the 20µl aliquots. The Dynabeads in the cells were manually removed using a DynaMag-2 Magnet (Thermo Fisher Scientific). 10µl counting beads was then added to each aliquot while sytox-AAD was added to the second aliquot only. The autofluorescence and cell count data were acquired by the same CytoFLEX S. 3000 counting bead events were collected for subsequent cell count analysis. Data acquired were analyzed using CytExpert and the cell number was calculated by following the manufacturer’s instruction.

### FACS analysis for T cell purity, activation markers and memory subsets

For activation markers and memory subsets analysis, cells were washed with PBS followed by staining with LiveDead-violet for 20 min at room temperature. Cells were subsequently washed with FACS buffer and stained by incubating with fluorochrome-conjugated antibodies as specified in the relevant gating strategies in supplementary figures for 20 min at room temperature. For purity analysis, cells stained with LiveDead-violet were washed with FACS buffer followed by staining with FITC anti-CD3 and APC anti-CD19 antibodies for 20 min at room temperature. After staining, cells were washed twice with FACS buffer before data acquisition and the data were acquired by the same CytoFLEX S. Data acquired for activation and memory subsets were analyzed with FlowJo software (TreeStar) and the purity were analyzed using CytExpert. Examples of gating strategies for the analysis of T cell activation and memory subsets are illustrated in Suppl. Figure 2 and 3.

### In vitro cytotoxicity assay

Cryopreserved NTC and CD19TDC were thawed and cocultured with NALM6-luciferase cells (NALM6 cells) at various effector (GFP^+^CD19TDC) to target cell (NALM6 cells) ratios (E:T) for 18 hours in RPMI1640 medium (Cytiva) supplemented with 10% FBS and 1% penicillin-streptomycin (Thermo Fisher Scientific). After co-culture, cells were transferred to a white 96 well opaque plate (SPL Life Sciences) followed by adding the Bright-Glo Luciferase Assay reagents to the wells (Promega). The luminescence signals were measured using Infinite M200Pro plate reader (Tecan).

### Analysis of metabolites in the spent cell-free culture supernatants

Spent culture supernatants collected were spun down at 500g, 5min for removing the cells. The supernatants were then kept in -80°C for use. The concentrations of glucose, lactate, glutamine and glutamate in the supernatants were measured by reagents: Glucose Bio, Lactate Bio, Glutamine V2 Bio and Glutamate V2 Bio, respectively using Cedex Bio Analyzer (Roche CustomBiotech).

### Statistical analysis

Data are expressed as the mean ± standard deviation (SD). Pearson correlation analysis is used for calculating the correlation and the goodness of fit is expressed as the coefficient of determination (r^2^). The best-fit line is determined by simple linear regression and the 95% confidence bands of the best-fit line is shown. Statistical significance was determined by tests as mentioned using GraphPad Prism software (GraphPad software). P values smaller than 0.05 were considered statistically significant. For significant results, the strength of significance is designated in terms of P values of *<0.05, **<0.01, and ***<0.001, ****<0.0001.

## Results

### NADH and FAD autofluorescence can be analyzed by 405nm laser equipped flow cytometer

Previous studies have demonstrated the UV-laser equipped flow cytometer can be used for measuring the change of autofluorescence in immune cells after T cell activation^17^. However, the use of UV laser is genotoxic for T cells, which is not suitable for intermittent monitoring of CAR-T cells in production. In order to investigate the cellular autofluorescence can be used for monitoring cell production, we investigated the use of a 405nm violet laser instead of a UV laser for detecting the changes in autofluorescence in T cells after activation. CD3^+^ T cells were activated with Immunocult and the changes in NADH and FAD autofluorescence signals were measured using a flow cytometer equipped with a 405nm laser. We observed that the percentages of NADH^+^ and FAD^+^ cells were significantly upregulated in a time dependent manner from day 1 to 3 after T cell activation when comparing against day 0 (Figure 1A). The mean fluorescence intensity (MFI) of the activated T cells for NADH and FAD significantly increased in the same manner (Figure 1A). To investigate whether the upregulation in NADH and FAD correlated with T cell activation, we measured the upregulation of the activation markers CD25 and CD69 in parallel. We showed that percentage and MFI of CD69 expression were increased after T cell activation (Figure 1B). The result is in line with a previous study which showing NADH and FAD are upregulated together with CD69 after 24 hours^16^. Although the percentage change of CD25 was not significant at early time points, the fold change of MFI of CD25^+^ cells increased significantly after day 1 post activation (Figure 1B), and were similar to those of NADH and FAD after T cell activation. As expected, the increase in percentage and MFI of NADH and FAD have a strong and significant positive correlation with the increase in the fold change of MFI of CD25^+^ cells (Suppl. Figure 4 and Figure 2). These results suggested that the 405nm violet laser can measure the changes of autofluorescence signals for both NADH and FAD in activated T cells, and that the upregulation of NADH and FAD were correlated with T cell activation.

**Fig. 1.**
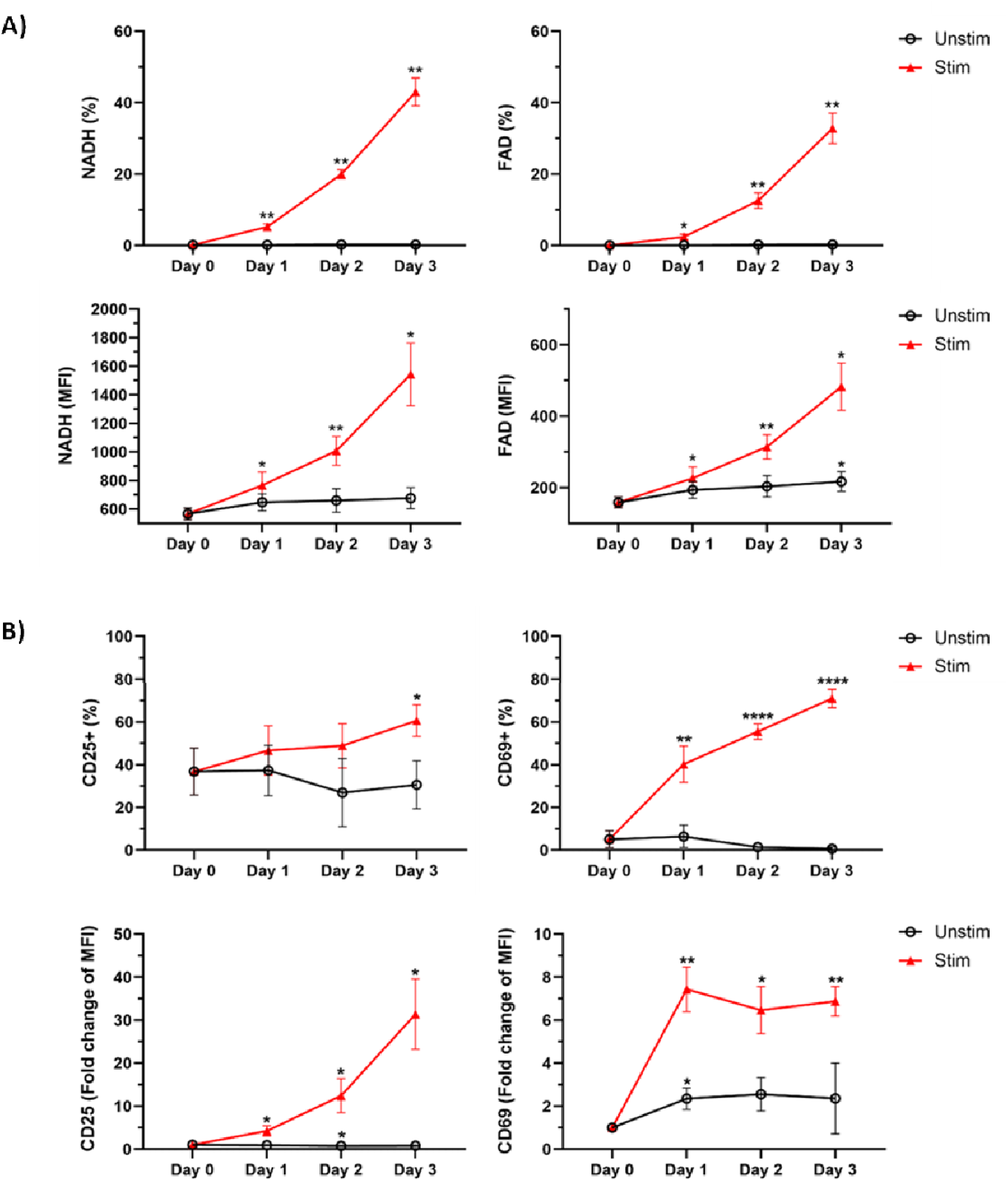
The increase of NADH and FAD autofluorescence after T cell activation can be measured by 405nm violet laser. CD3 T cells were stimulated with Immunocult for 3 days. NADH and FAD autofluorescence, CD25 and CD69 expression were measured at indicated time points by a 405nm violet laser equipped flow cytometer. A) Time dependent increase of the percentage of expression and mean fluorescence index (MFI) of NADH and FAD in T cells after stimulation (donors =3). B) The changes in the percentage and the fold changes of MFI of CD25 and CD69 expression in T cells after stimulation. The fold change of MFI is obtained by dividing the MFI of CD25^+^ or CD69^+^ cells by their counterparts on day 0 (donors =3). Unstimulated cells served as the negative controls for results in A) and B). Results for A) and B) are expressed as mean ± SD. Unpaired t test with Welch’s correction was used for statistical analysis. *P<0.05, **P<0.01, ****P<0.0001 compared against day 0 unstimulated cells.

**Fig. 2.**
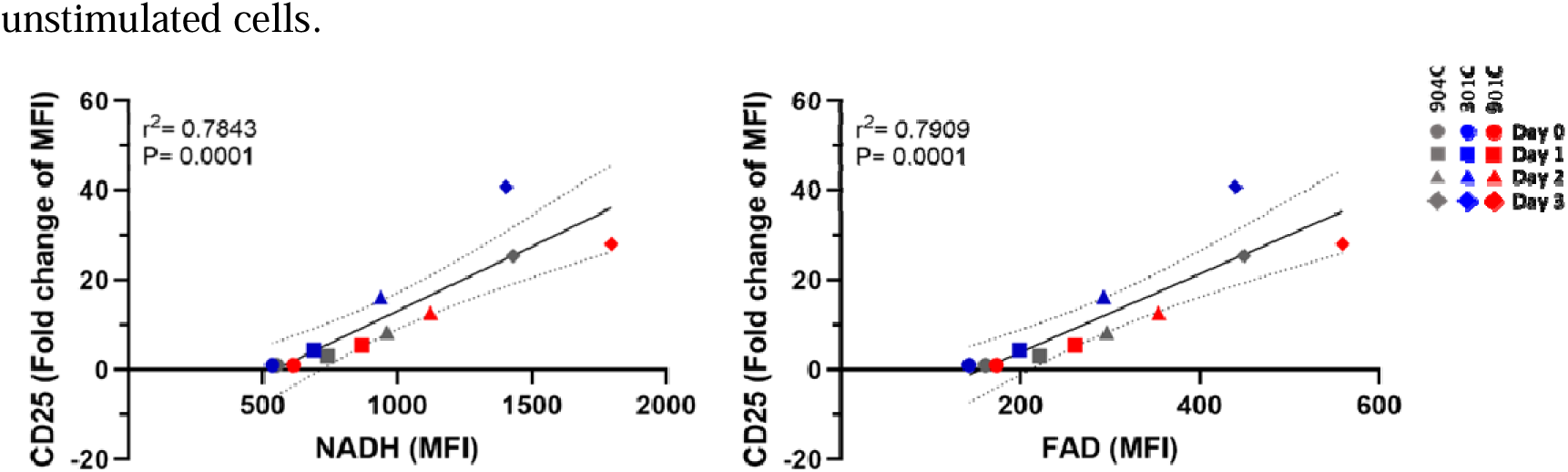
The increase of NADH and FAD autofluorescence after T cell activation significantly correlate with the expression of CD25. Results of Pearson correlation analysis show that the increase of the MFI of NADH and FAD of activated T cells are significantly correlated with the increase of the fold change of MFI of CD25^+^ activated T cells after stimulated by Immunocult (donors =3).

### NADH and FAD correlates with extracellular lactate in activated T cells

Upon T cell activation, T cells upregulate glycolysis and glutaminolysis for providing energy to support T cell proliferation and effector functions^23–26^. The increased consumption of glucose for glycolysis and glutamine for glutaminolysis along with the increased production of their respective metabolic products, lactate and glutamate can be reflected by measuring the changes of metabolites in the cell culture supernatant^21^. To do this, the spent media obtained from the unstimulated and stimulated T cells from day 1 to 3 were collected and subjected to metabolite analysis. We observed that the concentration of lactate was significantly increased from day 1 to day 2 and from day 2 to day 3, while a significant increase of glutamate concentration was only observed from day 2 to day 3 (Figure 3A). A significant reduction of glutamine could be observed from day 1 to day 2 and from day 2 to day 3, while only a slight but insignificant reduction was found for glucose (Figure 3A). A previous study revealed that extracellular lactate can be used for measuring T cell activation and proliferation^27^. Since no cell labeling step is required for measuring the changes of lactate in the spent media, lactate therefore can be a label-free parameter for measuring T cell activation. In order to investigate whether NADH and FAD can be an alternative label-free method which is comparable to lactate for measuring T cell activation, the correlation between extracellular lactate and NADH or FAD, both in percentage and MFI, from day 1 to 3 after T cell activation were investigated. Results in Suppl. Figure 5 and Figure 3B showed that the increase in percentage and MFI of NADH and FAD are significantly and positively correlated with the increase of extracellular lactate after T cell activation, respectively.

**Fig. 3.**
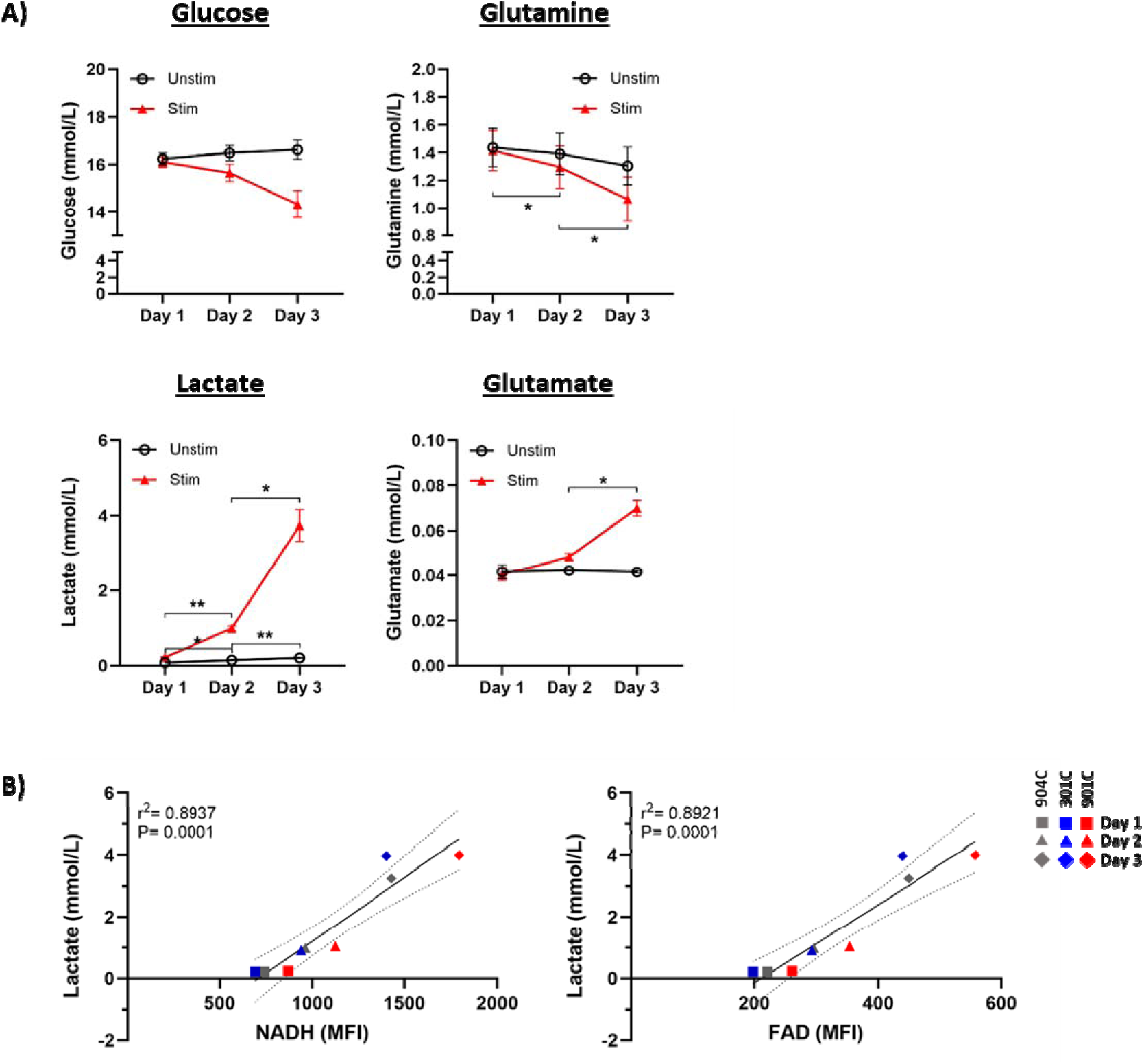
The changes in NADH and FAD autofluorescence is comparable to the changes of metabolites in the spent culture supernatants after T cell activation. A) CD3 T cells were stimulated with Immunocult for 3 days and the spent culture supernatants were collected at indicated time points for Cedex analysis. The glucose and glutamine concentrations are reduced along with the increase of lactate and glutamate concentrations in a time dependent manner after T cell activation (donors =3). Supernatants from unstimulated cells served as the negative controls. Results are expressed as mean ± SD. RM one way ANOVA with the Geisser-Greenhouse correction and Tukey’s multiple comparisons test, with individual variances computed for each comparison was used for statistical analysis. *P<0.05, **P<0.01 B) Results of Pearson correlation analysis show the increase of the MFI of NADH and FAD of activated T cells are significantly correlated with the increase of lactate concentration in the spent media after T cell stimulation by Immunocult (donors =3).

### Relationship between FAD autofluorescence, T cell proliferation and activation in a G-Rex bioreactor

As NADH and FAD autofluorescence can be used as a label-free cellular attribute for monitoring T cell activation, we aimed to understand the profiles of these two signals during T cell expansion to investigate whether they can be used as a cellular attribute for monitoring CAR-T production. T cells were cultured in a G-Rex bioreactor, and the changes of NADH and FAD in T cells were measured daily for 14 days after T cell activation to match current CAR-T cell production protocols^9,10^. After T cell activation, the percentage and MFI of NADH and FAD, similar to the above results, were increased for the first 3 days. The FAD signal peaked on day 4, followed by decreasing from ∼90% (90.4±4.8%) on day 4 to ∼14% (14.1±5.1%) on day 10, and subsequently became stable from day 10 to day 14 (13.9±4.6%). In contrast, the NADH signal stay elevated from day 4 (98.9±0.6%) to 6 (96.7±2.1%), then decreased from day 6 to ∼58% (57.8±14.0%) on day 12 (Figure 4A, left). On the other hand, the MFI of both NADH and FAD peaked at day 4 (NADH: 2958±166.7; FAD: 1155±66.9), followed by a continuous decrease until day 10 (NADH: 1380±65.2; FAD: 495.9±24.3), then became stable until day 14 (NADH: 1371±55.2; FAD: 500.2±21.5) (Figure 4A right). Since the autofluorescence from dead cells in the culture can affect the results, we also analyzed the NADH and FAD after excluding the dead cells using sytox-AAD, a dye for live and dead cells discrimination^17,28^. After excluding the dead cells, the NADH and FAD signals were comparable to the sytox-AAD unstained cells, demonstrating that dead cells did not meaningfully affect the interpretation of the results (Figure 4A). In summary, the changes of percentage or MFI of NADH and FAD are relatively consistent among different healthy donors over the 14 days culture period. Importantly, the changes of MFI of NADH and FAD are very similar as shown in figure 4A right. The similar MFI results of these two signals therefore implied that the MFI of either NADH or FAD can be used independently for monitoring T cell activation and expansion.

**Fig. 4.**
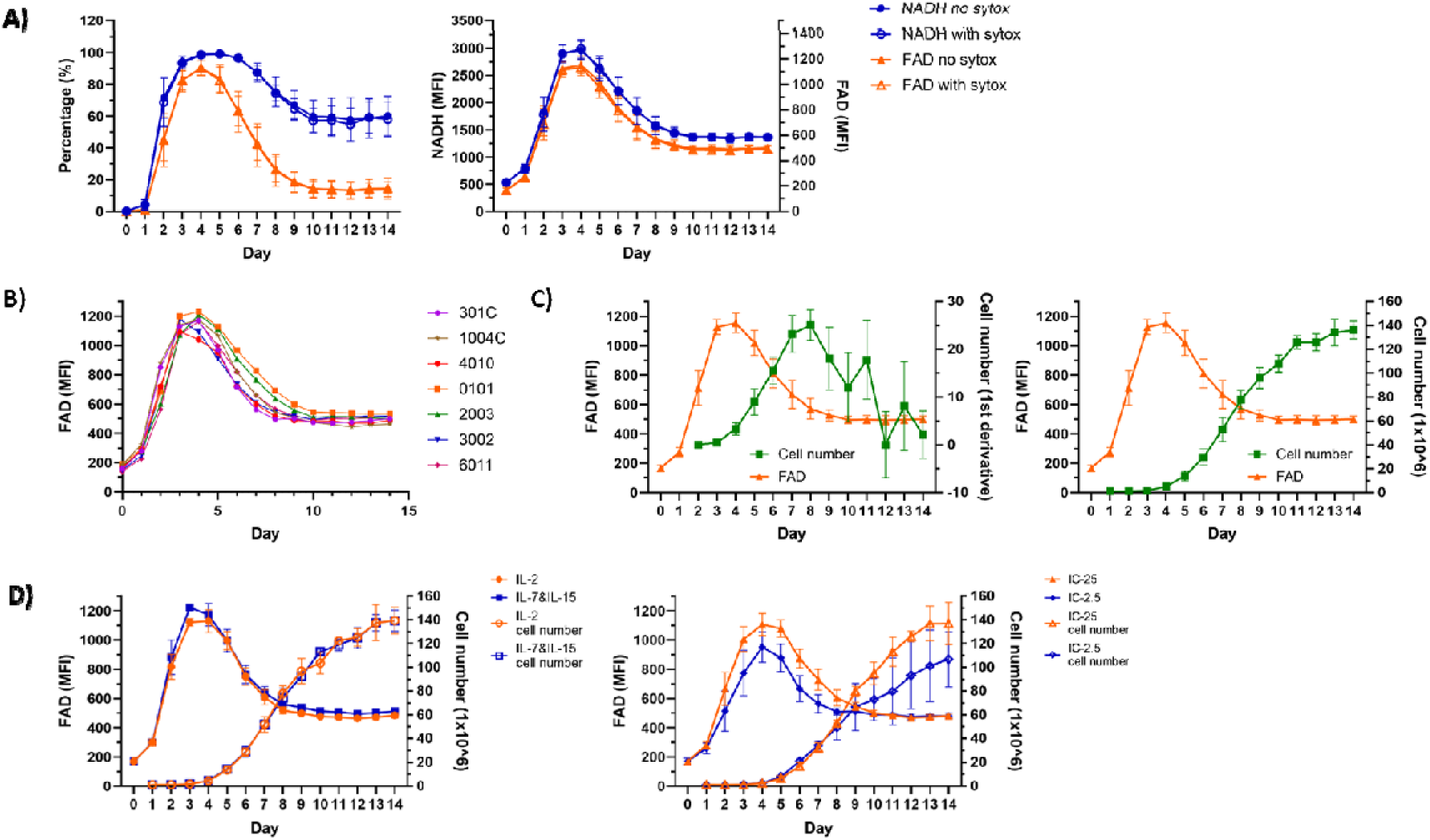
Longitudinal changes of FAD autofluorescence in activated T cells over 14 days. CD3 T cells were stimulated with Dynabeads or Immunocult (where specified) and the NADH and FAD autofluorescence were measured daily for 14 days. A) Comparable curve patterns between the unstained cells and the sytox-AAD stained dead cell excluded cells for the percentage of expression (left) and MFI (right) of NADH and FAD (donors=7). B) The longitudinal changes of the MFI of FAD in T cells of different individuals (donors =7). C) The longitudinal changes of the MFI of FAD in T cells versus the first derivatives of T cell number (left) and the T cell numbers in the culture (right) (donors =7). D) Comparisons of the MFI of FAD and T cell numbers between T cells activated by Dynabeads cultured in IL-2 alone or in a combination of IL-7 and IL-15 (left) (donors =3). Comparisons of the MFI of FAD and T cell numbers between T cells activated by 25µg/ml (IC-25) (donors =3) or 2.5µg/ml (IC-2.5) (donors =2) Immunocult (IC) (right). All results are expressed as mean ± SD.

In the subsequent experiments, we focused on the MFI of FAD for further study for two major reasons. First, FAD can be excited by a longer wavelength of light, which favors its future application in cell production monitoring^14^. Second, while assessments based upon percentage of positive cells can be subjective as it was achieved by manual gating, MFIs are obtained from the whole histogram, with no need for manual gating. When we analyzed seven healthy donors individually, we observed that while there were some differences in the day of peak and the rate of signal decrease, the changes of the MFI of FAD were similar (Figure 4B). We then examined the relationship between the changes of the MFI of FAD and the cell proliferation rate. The T cell proliferation rate started to increase on day 4, which is the peak of the MFI of FAD. The proliferation rate accelerated from day 5 to day 7, with a rapid decrease in the MFI of FAD, followed by a slowing at day 8 as the changes in the MFI of FAD flatten. (Figure 4C, left). The comparison between the MFI of FAD and absolute cell number is shown in Figure 4C, (right). These results suggested the MFI of FAD can be used to monitor the rate of T cell proliferation in the G-Rex platform.

Next, we evaluated the potential factors that could affect the MFI of FAD. We found that when the T cells were stimulated with same concentration of Dynabeads which implied same T cell stimulation strength, a change of cytokines from IL-2 to a combination of IL-7 and IL-15 had no impact on the MFI of FAD (Figure 4D, left). In contrast, when the cells were exposed to less Immunocult T cell activator, which reduces T cell stimulation strength, the peak MFI of FAD was lower and correlated with a lower value of the MFI of FAD from day 1 to day 4 when compared to its higher concentration counterpart (Figure 4D, right). This result is consistent with our results showing that the MFI of FAD correlated with the T cell activation (Figure 2). In summary, these results suggested that the MFI of FAD is affected by the strength of stimulation instead of the cytokines and can be applied to T cells stimulated by different activators.

### Using FAD autofluorescence as a signal for harvesting the CD19CAR-T cells (CD19TDC)

Since the changes of the MFI of FAD were similar among T cells obtained from different individuals and associated with the T cell proliferation rate, we investigated whether the MFI of FAD can be a cellular attribute for improving the CAR-T cell production process. By comparing the rate of change in the MFI of FAD and the rate of change in cell number, we found that the day after the minimum of the rate of decrease for the MFI of FAD (day 7) was the day before the proliferation rate started to decrease (day 8) (Figure 5A). These results suggested that the minimum of the 1^st^ derivative of the MFI of FAD can be used as a sign for harvesting the cells so as to maximize the cell number and at the same time, to avoid collecting the cells with reduced cell proliferation capacity. To evaluate whether this finding can be applied for CAR-T cell production, we transduced T cells with a lentiviral vector encoding a CD19 CAR one day after activation and monitored the changes in the MFI of FAD and cell number over 14 days. About 30 to 60% of T cells from the five different healthy donors tested expressed the CAR transgene after lentivirus transduction (Figure 5B). The bell-shaped curves for the MFI of FAD in NTC and CD19TDC are almost the same, suggesting that lentivirus transduction and CAR expression do not affect the FAD in CD19TDC (Figure 5C). Next, we compared the rate of change in the MFI of FAD and the rate of change in cell proliferation for CD19TDC. The rate of proliferation is slightly reduced on the day after the MFI derivative minimum (day 7), but is one day before the maximum turning point, which is the same as that in Figure 5A (Figure 5D). This result suggested that the minimum turning point of the MFI of FAD can be used as a signal for harvesting the CAR-T cells. Indeed, the fold change of cell number between days 7 and 10 only increases ∼2 (2.2±0.4) fold, as compared to a ∼10 (9.5±2.0) fold increase between day 4 to day 7, which is the day after minimum turning point. The fold change of cell number further reduced from day 10 to day 14, suggesting the CAR-T cells harvested on the day after the minimum turning point should have higher proliferation capacity than the cells harvested at the later time points (Figure 5E).

**Fig. 5.**
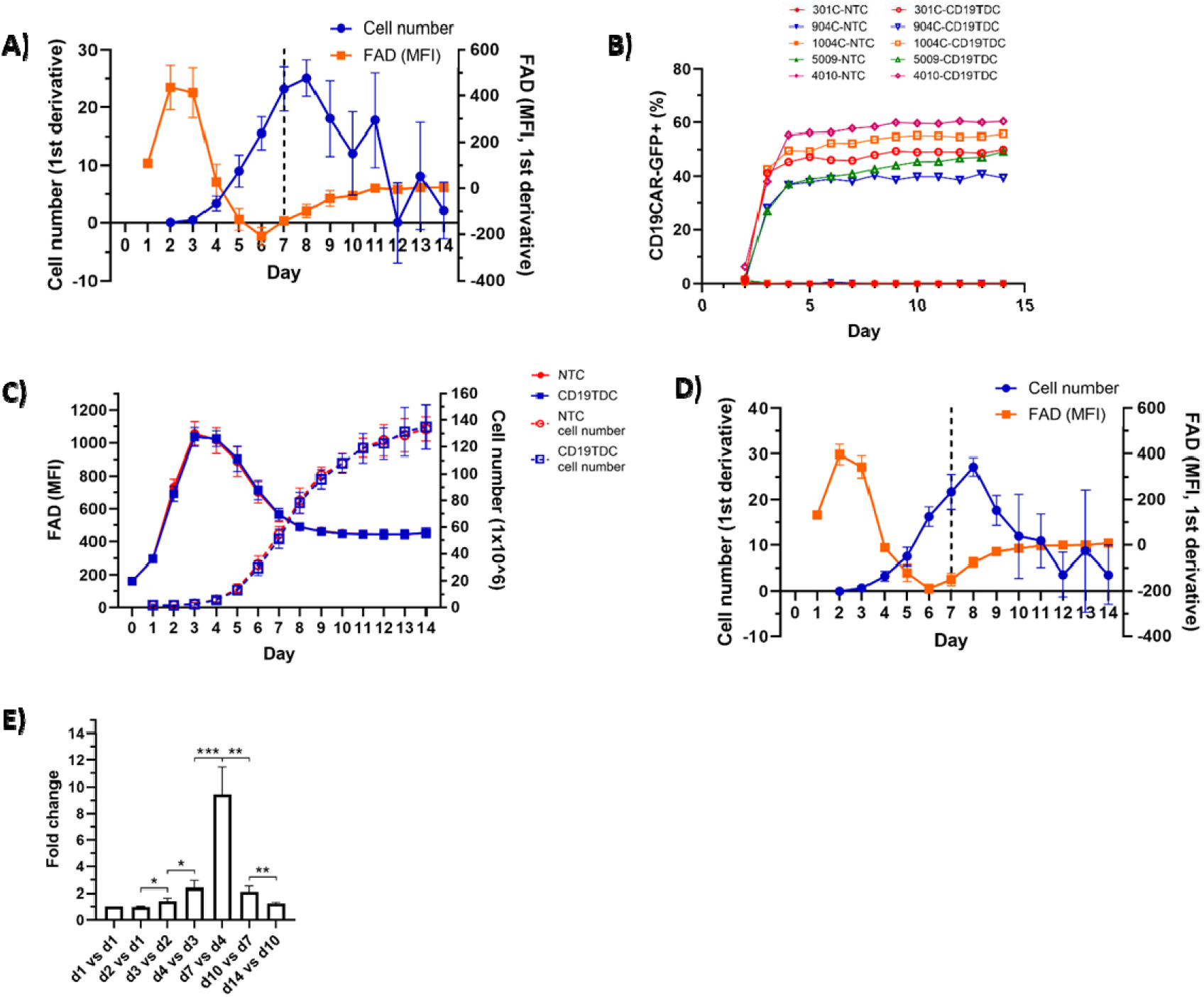
The first derivative of FAD autofluorescence reaches minimum and rebound before the decrease of proliferation rate in T cell and CD19CAR-T cells (CD19TDC). A) The first derivatives of the MFI of FAD of T cells reached minimum and rebound (indicated by dotted line) one day before the first derivatives of T cell numbers started to decrease (donors =7). B) CD19CAR expression on T cells after lentivirus transduction as determined by GFP expression. Non-transduced cells (NTC) served as the negative control for transduction (donors=5). C) The longitudinal changes of the MFI of FAD in NTC and CD19TDC versus their corresponding cell numbers (donors =5). D) The first derivatives of the MFI of FAD of CD19TDC reached the minimum and rebound (indicated by dotted line) one day before the first derivatives of the cell numbers of CD19TDC started to decrease (donors=5). E) The fold change of CD19TDC cell numbers between the indicated time points (donors=5). Results for A), C), D) and E) are expressed as mean ± SD.

### Comparisons of the memory subsets and cytotoxicity between the CD19TDC harvested on the day after minimum turning point of FAD autofluorescence and on day 14

The differences in the memory T cell phenotypes and the cytotoxic function of the CD19TDC harvested on the day after minimum turning point and on day 14 were investigated. Figure 6A shows the day after the minimum turning point for the 3 different donors tested are on day 7. While the differences are not significant, the cells collected on day 7 had slightly more naïve/stem cell memory T cells (T_N/SCM_) and central memory T cells (T_CM_), while the day 14 cells had more effector memory T cells (T_EM_) and terminally differentiated effector cells (T_EMRA_). This result, in line with previous studies^29,30^, suggested that T cells collected at earlier time points contain more less differentiated T cells (Figure 6B). Next, the cytotoxic function of the day 7 and day 14 CD19TDC were compared by co-culturing with the NALM6 cell line *in vitro*. Both day 7 and day 14 cells can kill the NALM6 cells at effector (E: CD19TDC) to target cell (T: NALM6) ratio of 1:2 and the percentage of specific lysis increased as the E:T ratio increased. Except donor 301C, the killing capacity of day 7 cells were always significantly lower than that of day 14 cells (Figure 6C). A possible reason of this can be because there were more T_EM_ and T_EMRA_, which are more cytotoxic than T_N/SCM_ and T_CM_ in the day 14 cells. The difference of donor 301C could be due to donor-to-donor variation. Thus, T cells that are harvested at the day after minimum turning point equipped with more less differentiated T_N/SCM_ and T_CM_ that are capable of killing the target tumor cells.

**Fig. 6.**
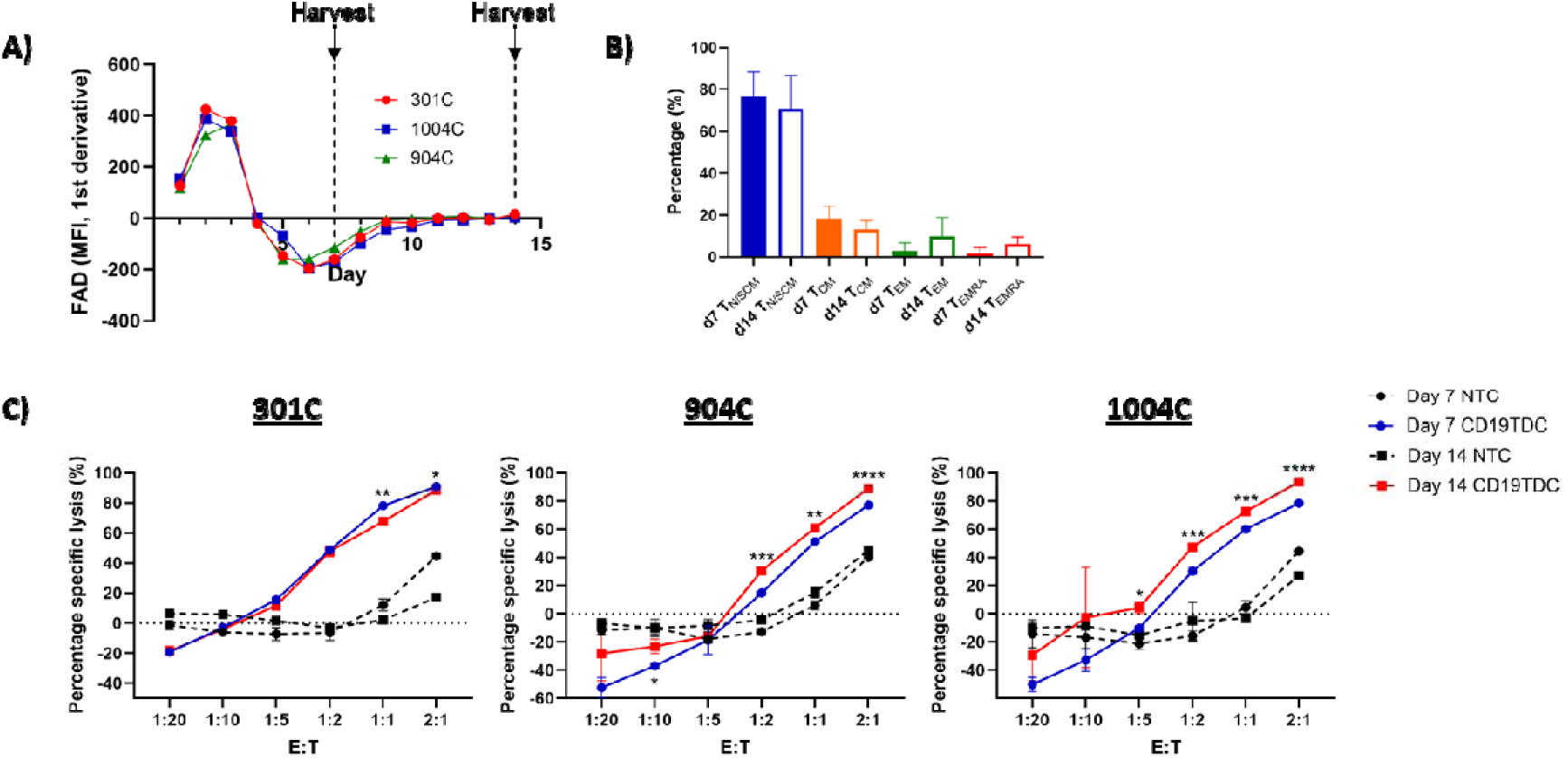
Using FAD autofluorescence as a CQA for CD19TDC production. CD19TDC were prepared for memory subsets analysis and cytotoxicity assay. A) The first derivatives of the MFI of FAD against time of 3 different donors. CD19TDC were harvested on day 7 when the first derivatives of the MFI of FAD reached the minimum and rebound (indicated by dotted line) and on day 14 which is the endpoint of the experiment. B) FACS analysis showed no significant differences for each memory T cell subset between day 7 and day 14 GFP^+^ CD19TDC (donors=3). C) The specific killing was performed by co-culturing the GFP^+^ cells of CD19TDC with NALM6-luc cells at effector to target cell ratio (E:T) as indicated. Results of 3 different donors showed day 7 and day 14 CD19TDC started to kill the target cells effectively since E:T=1:2 in a dose dependent manner. Results for B) are expressed as mean ± SD and C) are the mean ± SD of 3 technical replicate measurements for each donor. Unpaired t test with Welch’s correction was used for statistical analysis. *P<0.05, **P<0.01, ***P<0.001, ****P<0.0001

## Discussion

Due to the positive clinical outcome of CAR-T cell therapies in relapsed/refractory haematological malignancies, and the large number of new candidates currently undergoing clinical trials, the demand for engineered immune cell therapies is anticipated to increase in future^6,7^. Several major challenges for scaling up and scaling out the production are associated with the complex manufacturing process, limited capacity of manufacturing sites and the need for a highly skilled workforce^11,12^. Therefore, future advancements in CAR-T cell manufacture will focus on the development of closed and automated systems. Current commercial bioreactors such as CliniMACS Prodigy and Lonza Cocoon have made CAR-T cell manufacturing more automated, however, cell sampling during production still requires manual handling through the sterile tubing welders or aseptic access ports^12^. While safety measures have been implemented during manual sampling, the most effective way to minimize the opportunistic contamination is to avoid intervening with the cell culture. Hence, the development of label-free technologies for monitoring CAR-T cell manufacturing processes in closed and automated systems is essential. Previous studies have shown that NADH and FAD autofluorescence can serve as label-free biomarkers for T cell activation and their changes in T cells after activation can be analyzed using a flow cytometer equipped with a UV laser or two-photon microscopy^15–17^. Unfortunately, the genotoxicity of UV wavelengths, short imaging depth and a requirement for settled cells hinder the application of these technologies for screening autofluorescence changes in CAR-T cells during manufacture. Here, we demonstrated that a less harmful 405nm violet laser can be used for the rapid detection of the changes of NADH and FAD autofluorescence in T cells after activation and during expansion. This approach provides a potential method for leveraging autofluorescence to monitor CAR-T cell manufacture processes in closed and automated systems.

T cell activation, genetic modification, and expansion are critical stages for CAR-T cell manufacturing^3,8,9^. Since extracellular lactate in cell culture supernatants can be a label-free approach for assessing T cell activation and cell proliferation^27^, the significant positive correlations between NADH and FAD autofluorescence and the lactate concentration in the spent media, as well as the T cell activation marker CD25 in the first 3 days after T cell activation, demonstrate that NADH and FAD autofluorescence can serve as an alternative label-free approach for measuring T cell activation. Since NADH and FAD autofluorescence showed significant changes one day after T cell activation, this approach will be applicable for next-generation, shortened CAR-T cell manufacturing protocols^29,30^. As the changes in percentage and MFI of both NADH and FAD in T cells were comparable in T cell activation and expansion, we decided to focus on the MFI of FAD for our studies. This is because the maximum excitation wavelengths for FAD is not only longer than that for NADH, but also can extend beyond 405nm, thereby reducing potential damages to the cells^14^. Additionally, since the results expressed in MFI were obtained from the entire histogram, the results were not subjective when compared to those expressed in percentage which required manual gating. The use of visible lights with longer wavelengths than 405nm for FAD autofluorescence measurement deserves for further investigation to determine whether similar results can be obtained.

Previous studies revealed that CAR-T cells produced with shorter production protocols not only retain the ability to kill their respective target cells *in vitro*, but also exhibit enhanced tumor-killing efficacy *in vivo*^29,30^. Our findings demonstrated that the killing capacity of CAR-T cells harvested on the day after the MFI derivative minimum of FAD was comparable to those harvested on day 14 cells *in vitro*. Further experiments will be required to test on patient samples, as well as to investigate the persistence and the tumor-killing function of these CAR-T cells *in vivo* so as to validate the use of the minimum turning point of the rate of change in MFI of FAD as a signal to shorten the manufacturing process. In contrast to the conventional CAR-T cell therapies which utilize autologous T cells from patients, allogeneic CAR-T cells derived from T cells of healthy donors can possibly be developed into “off-the-shelf” products in the future^31,32^. Our results from healthy donor-derived cells reveal that the minimum turning point of the rate of change in the MFI of FAD can potentially be leveraged in improving the manufacturing process by maximizing the yield of allogeneic CAR-T cells in a shorter time while preserving their proliferative capacity.

## Conclusions

In conclusion, we have established a method to use 405nm visible light to measure the NADH and FAD autofluorescence for investigating the T cell activation and expansion. Also, we identified that the minimum turning point of the rate of change in MFI of FAD can be a potential signal indicating how to optimally time CAR-T cell harvest. Since this measurement relies upon visible light, it would be feasible to incorporate autofluorescence measurements for FAD into closed and automated systems for real time monitoring of CAR-T cell manufacturing.

## Author contributions

K.-W.C and M.E.B. conceived the project and planned the experiments. K.W.C., W.-X.S, F.K., D.B.L.T., conducted the experiments and collected the data. K.-W.C., analyzed the data. Y.H. Lee and M.E.B supervised the work. K.-W.C. and M.E.B wrote the manuscript. All authors reviewed and approved the manuscript.

