## Supplementary Materials for "Use of cellular FAD autofluorescence as a label-free cellular attribute for the production of chimeric antigen receptor-T cells"

### Supplementary figures

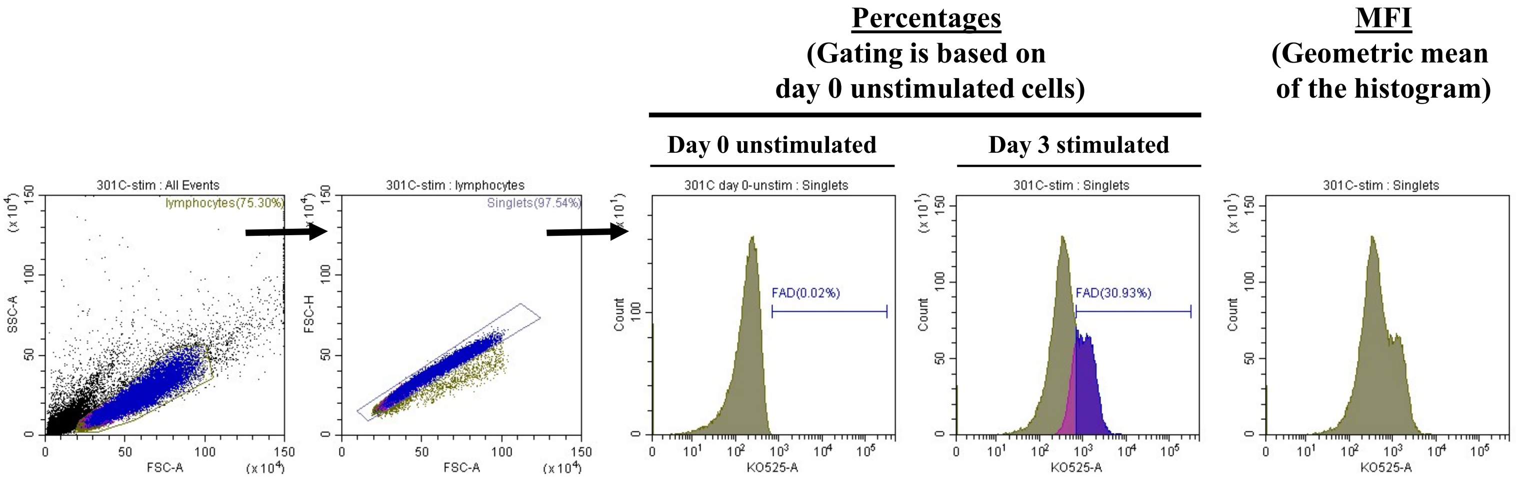


**Suppl. Fig.1. Gating strategy for measuring NADH and FAD autofluorescence in unstained T cells.**
Using the analysis for FAD autofluorescence as an example. T cells were gated followed by gating on the singlets for analysis. The gates for measuring the percentage of NADH and FAD expression were set based on the unstimulated cells on day 0. The mean fluorescence intensity (MFI) of NADH and FAD is the Geometric mean of the corresponding histogram.

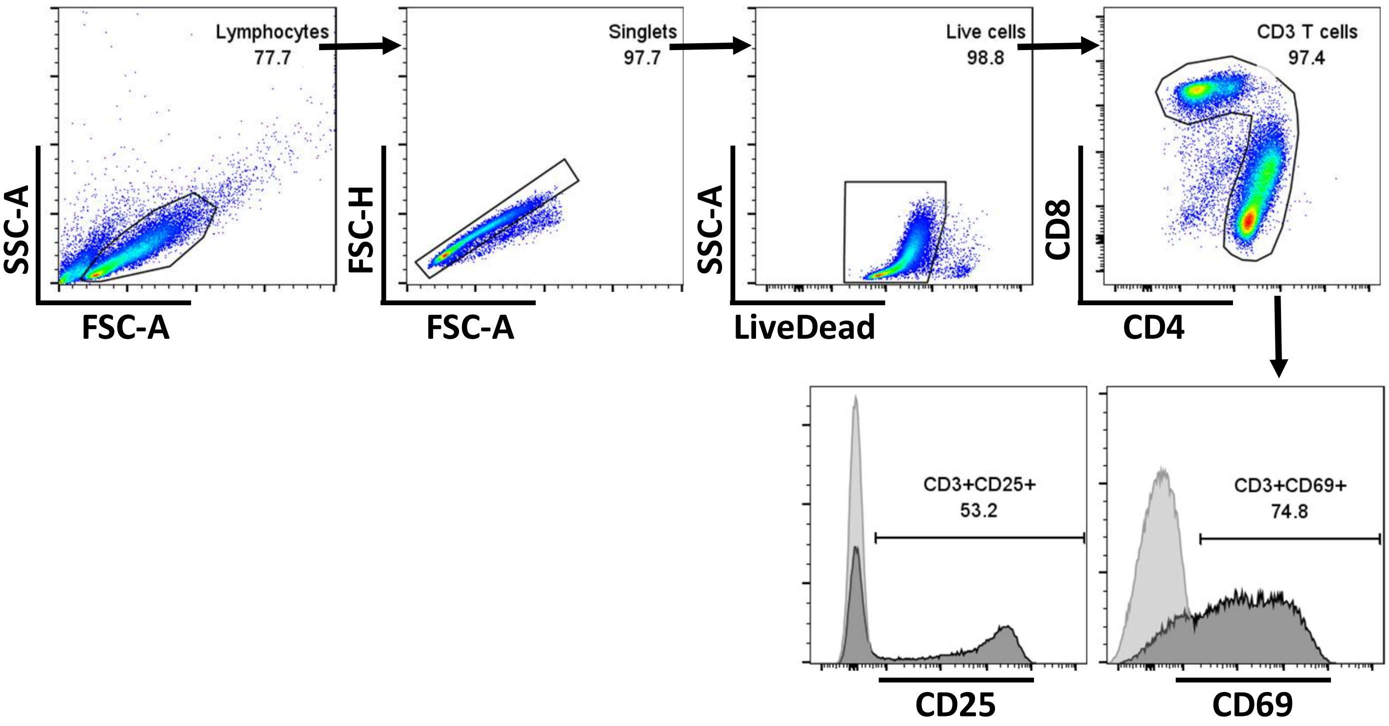


**Suppl. Fig.2. Gating strategy for measuring the CD25 and CD69 expression of T cells.**

T cells were gated followed by gating on the singlets. The dead cells were excluded by gating on the LiveDead-violet negative cells. The expression of CD25 and CD69 on CD3 T cells were analyzed by gating on the CD4 and CD8 T cells together.

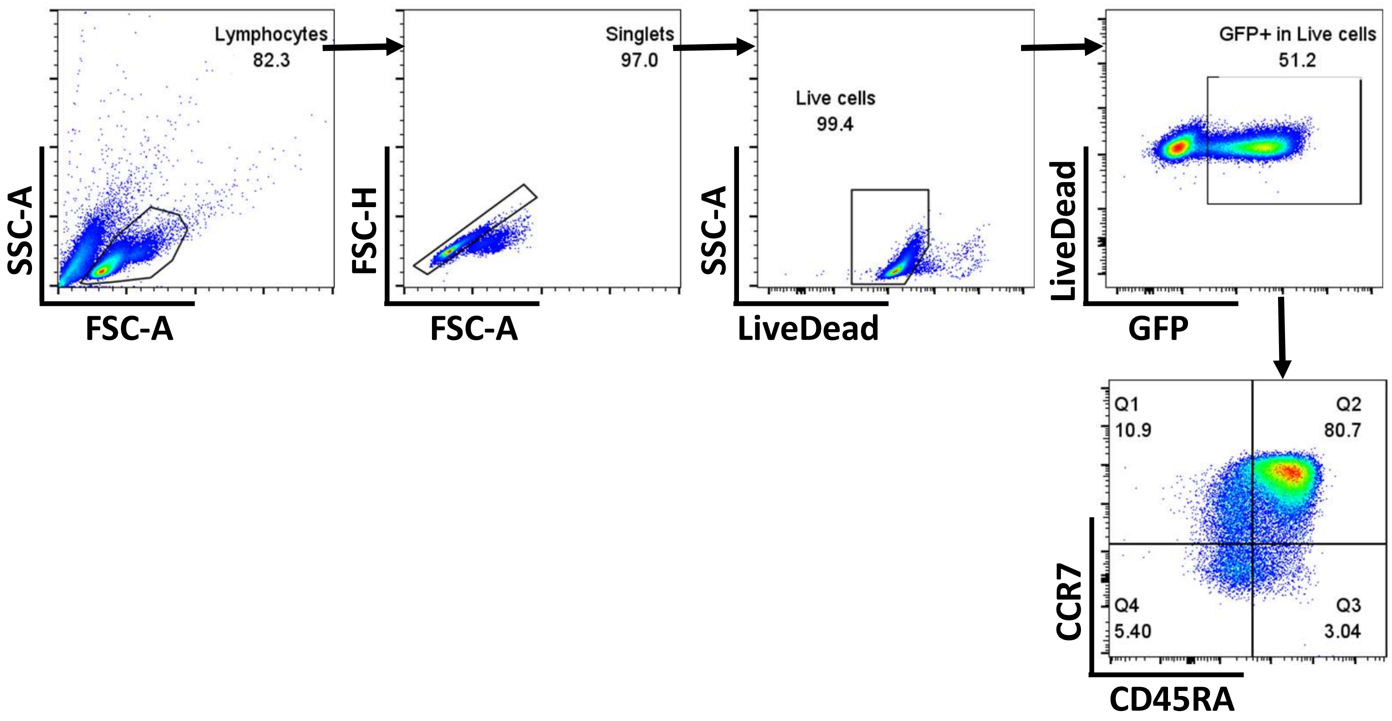


**Suppl. Fig.3. Gating strategy for investigating the memory T cell subsets in CD19CAR expressing T cells.**
T cells were gated followed by gating on the singlets. The dead cells were excluded by gating on the LiveDead-violet negative cells. The $GFP^{+}$ CD19CAR expressing cells were then gated for memory subsets analysis.

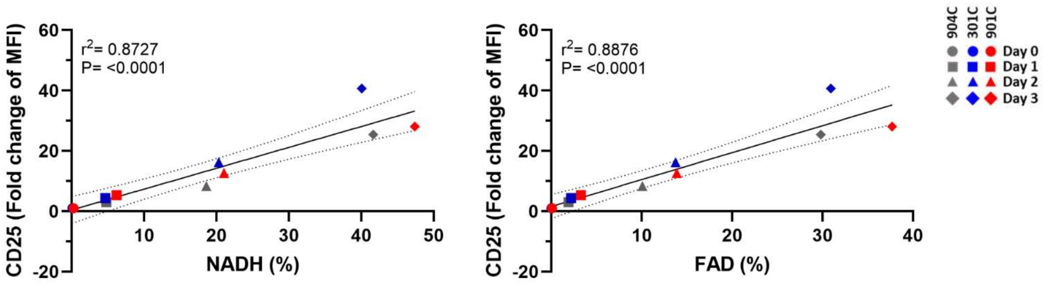


**Suppl. Fig.4. The increase of NADH and FAD autofluorescence after T cell activation significantly correlate with the expression of CD25.**
Results of Pearson correlation analysis show that the increase of the percentage of $NADH^+$ and $FAD^+$ cells are significantly correlated with the increase of the fold change of MFI of $CD25^+$ activated T cells after stimulated by Immunocult (donors =3).

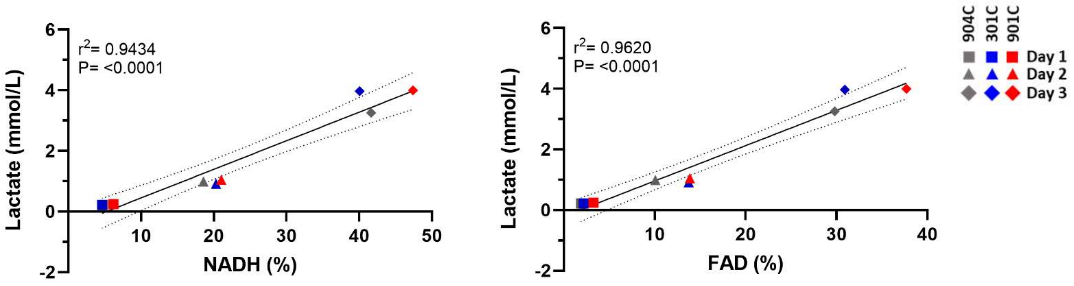


**Suppl. Fig.5. The increase of the percentage of NADH and FAD autofluorescence is positively correlated with the increase of lactate concentration in activated T cells.**

Results of Pearson correlation analysis show that the increase of the percentage of $NADH^+$ and $FAD^+$ cells are significantly correlated with the increase of lactate concentration in the spent media after T cell stimulation by Immunocult (donors =3).
